# Integrating MHC Class I visibility targets into the ProteinMPNN protein design process

**DOI:** 10.1101/2024.06.04.597365

**Authors:** Hans-Christof Gasser, Diego A. Oyarzún, Javier Alfaro, Ajitha Rajan

**Affiliations:** School of Informatics, University of Edinburgh, Edinburgh, UK; School of Biological Sciences, University of Edinburgh, Edinburgh, UK; International Centre for Cancer Vaccine Science, University of Gdańsk, Gdańsk, Poland; Department of Biochemistry and Microbiology, University of Victoria, Victoria, Canada; The Canadian Association for Responsible AI in Medicine, Victoria, Canada

**Keywords:** Deimmunization, ProteinMPNN, MHC Class I, Machine Learning, Language Models

## Abstract

`ProteinMPNN` is crucial in many protein design pipelines, identifying amino acid (AA) sequences that fold into given 3D protein backbone structures. We explore `ProteinMPNN` in the context of designing therapeutic proteins that need to avoid triggering unwanted immune reactions. More specifically, we focus on intra-cellular proteins that face the challenge of evading detection by Cytotoxic T-lymphocytes (CTLs) that detect their presence via the MHC Class I (MHC-I) pathway. To reduce visibility of the designed proteins to this immune-system component, we develop a framework that uses the large language model (LLM) tuning method, Direct Preference Optimization (DPO), to guide `ProteinMPNN` in minimizing the number of predicted MHC-I epitopes in its designs. Our goal is to design proteins with low MHC-I immune-visibility while preserving the original structure and function. For our assessment, we first use AlphaFold to predict the 3D structures of designed protein sequences. We then use TM-score, that measures the structural alignment between the predicted design and original protein, to evaluate fidelity to the original protein structure. We find our LLM-based tuning method for constraining MHC-I visibility is able to effectively reduce visibility without compromising structural similarity to the original protein.

## 1 Introduction

The popular protein design tool `ProteinMPNN` [1] tackles the task of predicting AA sequences given a desired 3D protein backbone structure (backbone). These backbones are often generated by other models - like the diffusion based Rfdiffusion [2] and FoldingDiff [3].

Some of the proteins designed using `ProteinMPNN` might be therapeutic proteins, in which case the design process will have to take into account the reaction of the immune-system to these non-self proteins. The consequences of disregarding the immune reaction could range from reduced therapeutic efficacy up to unwanted auto-immune reactions. These could target the therapeutic protein itself - as demonstrated by anti-transgene immunity [4, 5]. However, also other components of the medicine like the CRISPR-Cas9 gene editing machinery [6] or the viral vector [7, 8] could potentially attract unwanted attention by the immune-system.

The immune-system components that need to be taken into consideration will primarily depend on the way the therapeutic is administered and the location, where the therapeutic is supposed to unfold its effect. For extracellular proteins antibody (Ab) recognition and presentation by the MHC Class II (MHC-II) pathway will be most relevant. On the other hand, some proteins might have to be expressed within human cells - for example ribonucleic acid (RNA) therapeutics aimed at mitigating an issue within the cell. In these cases the MHC-I pathway is paramount.

In this paper we focus only on visibility to CTLs via the MHC-I pathway (see 2.1). To integrate this into the design process we tuned `ProteinMPNN` [1] (see 2.2) using DPO introduced in [9]. DPO adjusts the base network’s weights to increase the probability of generating sequences that better conform to our preferences (see 2.3). Our contribution is to apply this tuning technique to `ProteinMPNN`, generating sequences with reduced immune visibility and analyzing the trade-off between visibility reduction and sequence quality. The result is new models called `CAPE-MPNN`. Since they share the same architecture as `ProteinMPNN`, the `CAPE-MPNN` model weights can easily replace the original `ProteinMPNN` weights in existing protein design workflows.

## 2 Background

Protein folding prediction is concerned with finding the 3D structure that an AA chain will fold into. After significant advances in this area - in particular by models like *AlphaFold* [10, 11] - the inverse problem, protein design, has also attracted renewed attention. Traditional approaches have been physics based like the *Rosetta Packer* [12, 13]. The current machine learning (ML) revolution is leading to a plethora of data-driven approaches using for example LLMs [14, 15], Variational Autoencoders (VAEs) [16] and Generative Adversarial Networks (GANs) [17] to generate new amino-acid sequences. A particularly prominent model in this space is `ProteinMPNN` [1].

This can be embedded into various protein design workflows. A typical strategy might start by the generation of a backbone with a diffusion model like RFdiffusion [2] or FoldingDiff [3]. The 3D coordinates generated by these techniques can then be used as input to `ProteinMPNN` to generate AA sequence candidates. Another workflow might start with a known protein sequence for which we want to generate other sequences with similar function but different sequence. Deimmunization, where the goal is a protein that is less immune-reactive but functions similarly, motivates this application. In these cases at first the structure of the protein would be determined (experimentally or predicted) and this then used as input into `ProteinMPNN`.

Finally, the typical role of the ML step in the protein design process is to propose a set of candidate protein designs. These are usually assessed by a series of bioinformatics filters that rank them according to the likelihood that they will fold, function and conform to our preferences. This assessment might be computationally very costly, for example if Molecular Dynamics (MD) simulations are involved. The ML step should therefore design a set of candidates that is already enriched in a desirable property, and stand a good chance to pass these downstream filters. Due to its important role in the design of practically useful therapeutics, this project focus on reducing visibility to the immune-system.

### 2.1 MHC Class I pathway

There is a complex interplay between the components of the immune-system. However, for intra-cellular proteins the MHC-I pathway is of particular importance. This pathway is responsible for relaying information about what happens within a cell to surveying CTLs. These can destroy cells that they deem pathogenic.

To allow the immune-system to monitor cells’ health, the MHC-I pathway has evolved to present small sub-sequences (so called peptides, 8 to 10 AA long) of all proteins expressed within a cell on its surface. Here they can be surveilled by patrolling CTLs. In case an activated CTL detects a non-self peptide (a sub-sequence not encoded in that person’s genome), it will cause the destruction of the presenting cell.

In particular RNA therapeutics will be expressed within cells and their products therefore presented on the cell’s surface, highlighting the importance of incorporating MHC-I visibility into the protein design process for therapeutics. To integrate this into a particularly prominent protein design model, we explore the usage of DPO to tune `ProteinMPNN` to produce sequences with reduced visibility to CTL.

### 2.2 ProteinMPNN

Our tuned model `CAPE-MPNN` is based on `ProteinMPNN` [1] developed by Dauparas et al. `ProteinMPNN`’s basic functionality is to map a protein’s backbone geometry into AAs sequences that are predicted to fold into this given geometry. It supports protein complexes consisting of multiple chains. Another feature is that parts of the produced protein sequence can be fixed upfront. This might be particularly interesting when the user has detailed knowledge of the desired functional mechanism and wants to ensure the necessary AAs are present. Another quite useful feature is that they provide a score expressing the uncertainty of the design. They have trained various versions of this using different hyper-parameters. We present here the version we used to tune `CAPE-MPNN`. This uses the hyper-parameters listed in **Table 1**. The following description of `ProteinMPNN`’s architecture leaves out normalization, dropout and activation steps.

**Table 1:** ProteinMPNN hyper-parameters.

| Hyper-parameter | Symbol | Value |
| --- | --- | --- |
| Vocabulary size | V | 21 (20 standard AA + 'X') |
| Feature dimension | D | 128 |
| Nearest neighbors | N | 48 |
| Maximum number of residues in a batch | $T_{max}$ | 10,000 |
| Maximum sequence length | $S_{max}$ | 10,000 |

#### 2.2.1 Data

Tuning a network using DPO requires a preference dataset. This is made up of examples consisting of a prompt (a 3D backbone structure in our case), two possible responses generated by the model as well as the information which of those responses we prefer 2.3. We use the 3D backbone structures in the original `ProteinMPNN` dataset to produce this preference dataset. Therefore, we want to describe the original dataset here in a little bit more detail.

##### PDB Dataset

Their data examples are based on Protein Data Bank crystal structures with a resolution cutoff of 3.5 Å. Their initial dataset consists of 473,062 AA chains (106,344 unique AA sequences). PDB chains were clustered at 30% sequence identity using mmseqs2 [18] - so each cluster consists of similar chain-sequences. Each example in this PDB-Dataset is linked to a cluster. These were split into 23,349 training, 1,464 validation and 1,539 test clusters. When an example is requested, first a chain is sampled from this cluster using a uniform distribution. This chain is linked to a single PDB entry and the biological assemblies that the chain is part of within this entry are identified. If it is not part of any, then the chain alone is used. If it is part of several, a random biological assembly is picked. Then all the chains in the selected biological assembly are loaded and their coordinates transformed via the biological assembly specific transformations (chains can be in different positions and/or orientations in different biological assemblies). Many proteins consist of similar chains (e.g. homo-dimers). To prevent leakage of structural information from one chain to the other during training, they enforce that all chains with a pairwise TM-score above 0.7 with respect to the selected chain are also masked (no sequence information is provided) and have to be designed by the model. For the purpose of generating the preference data, we anyways always require the model to design all chains in an assembly.

##### Structure Dataset

The PDB Dataset described above is then used every two epochs to generate a Structure Dataset. This will have less examples, since in this step the maximum sequence length of the biological assemblies is restricted to 10,000 AAs (next to other drop-outs, *<* 10% shrinkage total). In this step they also remove leading or terminating histidine repeats. The batches supplied to the model are yielded from this dataset. A single batch has a maximum of 10,000 residues. This can be filled with a single very long biological assembly, or several smaller ones. Each of these sequences can comprise of multiple chains. We will refer to the number of sequences in a batch as *B* and the maximum length of any of those sequences as *S*. It holds *B · S < T*_*max*_.

#### 2.2.2 Embeddings

The examples delivered by the above dataset are processed within the model to produce the following initial embedding tensors that are then fed through the encoder and decoder which are updating these.

##### Edge embeddings

The edge embeddings tensor will have shape [*B, S, N, D*] (see **Table 1**). The dimensions two and three determine which residue and neighbor the encoding in dimension four belongs to. Only during `ProteinMPNN` training, Normal random noise with standard variation of 0.2 Å is added to the coordinates of each residue. The nearest *N* neighbors (including itself) are then determined based on *C*_*α*_ distance between residues. Then the coordinates of the AA atoms are used to calculate inter atom distances. For each residue-neighbor pair, 25 inter atom distances are calculated. Each of these is between an atom (N, C_*α*_, C, O and virtual C_*β*_) of the residue and an atom of the nearest neighbor (N, C_*α*_, C, O and virtual C_*β*_). Distances are not encoded as a single numbers, but rather using 16 radial basis functions (RBFs) with centers spread between 2 and 22 Å(far distances - above 25 Å-will have values close to zero in all bins). So, at this stage each residue-neighbor edge will be encoded using 400 elements (25 × 16). These get concatenated with a relative positional embedding. This embedding can be one of 66 different learnt 16 element vectors. One of these vectors signifies that the neighbor is on a different AA chain, while the others signify that the neighbor is up to 32 positions before or after the residue concerned (further distances are clipped).

Overall the concatenated edge embedding tensor now has a shape of [*B, S, N*, 416]. This is reduced to [*B, S, N, D*] by an affine transformation.

##### Node embeddings

These are a tensor of shape [*B, S, D*]. They start out being zero and are first updated by the encoder and afterwards by the decoder.

##### Token embeddings

These are only accessible by the *Sequence Decoder*. A sequence is represented as a list of AA indices (e.g. 0 is alanine, 1 is cysteine, …). These are embedded into a *D*-dimensional space resulting in a token embeddings tensor of shape [*B, S, D*].

#### 2.2.3 Architecture

`ProteinMPNN` is a relatively small model. The overall architecture has around 1.7m parameters. The subdivision Dauparas et al. make between a *Backbone Encoder* and *Sequence Decoder* is a useful way to think about the architecture. The encoder updates the initial edge and node embeddings. The decoder then predicts the AA sequence in an auto-regressive (AR) fashion. This means that, given the sequence created up to this point, the model produces a probability distribution over all possible AAs for the next AA in the sequence. While the standard procedure to train text AR models would be to use the standard left to right order within text, `ProteinMPNN` was trained to generate sequences in an arbitrary order (e.g. start with position 10, then sample position 50, 5, 23, …). For this the decoder can access the edge, node and token (of residues predicted so far) embeddings. The training objective was to minimize categorical cross-entropy per predicted AA sequence token.

We now delve into the information flow in a training forward pass. A sample step is the same for the encoder but the decoder needs to be called several times. This is a bit more complicated as it needs to be ensured that the decoder does not access information that is not yet available.

##### Backbone Encoder

We now will discuss the *Backbone Encoder*. This is a message passing neural network (MPNN) consisting of a stack of three encoder layers. It receives the edge embedding, node embedding, nearest neighbor information as well as a mask that stores which residues have no known coordinates. The edge and node embeddings are updated by each layer of the stack.

Within a encoder layer, at first a message tensor is produced. This happens by combining the edge and node embeddings into a tensor of shape [*B, S, N*, 3 *× D*]. For an edge between a residue specified in dimension 2 and one of its neighbors specified in dimension 3, its last dimension holds the following three embeddings (each *D* elements): the residue’s node embedding, the edge embedding, the neighbor’s node embedding. A sequence of three affine operations transforms this into the message tensor of shape [*B, S, N, D*].

Elements in the message tensor that are linked to missing residues are first masked out, before summing over its third (neighbor) dimension. The result of this is then added to the node embeddings (skip connection). These updated node embedding are then fed through a two layer fully connected network and the result added to the node embeddings again (skip connection).

To update the edge features, again a message tensor is produced as described above (using different parameters for the operations). This time, however, missing elements are not masked out. This message tensor is then added to the edge features (skip connection).

##### Sequence Decoder

A decoding order is sampled (missing residues are first). Based on this a backward and a forward mask are generated. An element in the backward mask is one if and only if a node’s neighbor has already been decoded. The opposite holds for the forward mask.

The decoder receives the finally updated edge and node embeddings from the encoder as well as the masked sequence embeddings. With those the tensor *EXV* of shape [*B, S, N*, 3 *× D*] gets produced. Its last dimension hold first the encoder produced edge embeddings, then zeros and finally the neighbor’s node embeddings (each *D* elements). In addition, the forward mask is applied to it (elements of neighbors that have already been decoded are set to zero).

Before the call of each of the three decoder layers, also another tensor *ESV* with a shape of [*B, S, N*, 3 *× D*] gets produced. Its last dimension holds initially the same elements at the beginning and end as *EXV*. Just the zeros in the middle are replaced by the AA embeddings of the neighbor. Also, the neighbor’s node embeddings at the end get updated with every decoder layer call. Finally, the backward mask is applied to *ESV* (elements of neighbors that have not yet been decoded are set to zero).

Each decoder layer is then called with the node embeddings (by reference), and the sum of *EXV* and *ESV*. So the following embeddings are available for each edge.

- The residue’s node embedding
- The edge embedding produced by the encoder
- The neighbor’s AA embedding or zero if the neighbor has not yet been decoded
- The neighbor’s node embeddings (encoder produced for not yet decoded neighbors)

The shape [*B, S, N*, 4 *× D*] tensor holding this per edge information is fed through three affine layers, resulting in a message tensor of shape [*B, S, N, D*]. The neighbor dimension of this message tensor is summed over and the result added to the node embeddings (skip connection). This updated node tensor is then fed through a two layer fully connected network and added to itself (skip connection). After this all node embeddings linked with unavailable residues are set to zero. After the last decoder layer, there is an affine transformation that produces logits from the final node embeddings.

### 2.3 Direct Preference Optimization

This technique is typically applied to LLMs to make them conform better to our preferences with regards to harmlessness and usefulness. It is derived from the more general method of reinforcement learning from human feedback (RLHF) [19] which was also used to tune the GPT foundation models to produce ChatGPT.

In a first step a foundation model (like OpenAI’s GPT-3) would be trained using self-supervised learning (e.g. on a corpus of text from the internet). Next word prediction is a common training objective. This foundation model will then be able to generate likely text, which is, however, not focused on a particular task - like answering questions, helping write text, summarizing text, … Therefore, typically, task specific data is then used to fine-tune the model for these, instead of just generating any kind of text. However, after this step, there typically can still be a lot of dangerous language left. To tune the model further, RLHF was introduced.

RLHF is based on examples generated by the fine-tuned foundation model. These examples are then rated by humans. A typical way to rate them is to give the human two generated texts and ask them to judge which one is better. This is the variant that DPO is based on. Alternatively, the human could also assign ratings. Independent of the concrete procedure, a reward model can then be trained in a supervised way based on this human produced preference data. This reward model predicts higher rewards for responses that are predicted to be preferred by humans. RLHF then relies on a reinforcement learning (RL) training step (typically using Proximal Policy Optimization (PPO) [20]) for which the reward model provides the rewards.

Tuning a model using RLHF can be quite difficult in practice. According to some practitioners, in particular selecting the many hyperparameters involved can be challenging. Therefore, a more straight-forward way to incorporate preferences into LLMs had to be found. For the RLHF version in which the feedback consists of humans choosing their preferred response from two generated ones, Rafailov et al. discovered such a more direct way. In [9] they show that this optimizes for the same objective as a supervised training way they called DPO.

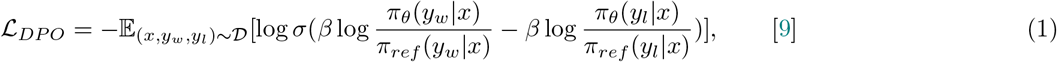

The training objective under DPO is to minimize the loss in **Equation 1**. There, *x* is the prompt for the generation (in our case the backbone we want to generate a sequence for). *y*_*w*_ is the preferred response, while *y*_*l*_ is the less preferred one. *π*_*ref*_ denotes the density function of the original model and *π*_*θ*_ that of the tuned model. An important parameter is *β*, which controls how far we want the model to steer from the original one. The higher *β*, the closer the tuned model will be to the original one.

## 3 Methods

**Box 1:**

**Terms and concepts**

- **MHC-I immune-visibility (or simply ‘immune-visibility’ or ‘visibility’):** The focus of our study is the MHC-I pathway which CTL utilize to detect destruction targets. For this study, visibility means the predicted number of 8, 9 and 10-mers within a protein sequence to be presented on the cell’s surface (overlaps counted separately). If a k-mer is presented by several MHC-I alleles this increases the visibility number only by one. A limitation of this approach is that, while visibility of a peptide to CTLs is a necessary condition for an immune-reaction it is not a sufficient one.
- **Visibility profile:** Different applications will require different interactions with the immune system. While it might be beneficial for vaccines to elicit an immune response to certain epitopes, therapeutic proteins will in general benefit from as little visibility as possible. With visibility profile we mean the conceptually desired visibility to the immune system. In this paper, we restrict ourselves to reducing the MHC-I visibility of designed proteins.

This section explains how `ProteinMPNN` was aligned with our immuno-visibility target using DPO. Afterwards, the metrics utilized to compare the various model checkpoints are introduced. Eventually, the DPO hyper-parameter search is outlined.

### 3.1 Data

We use two sources of data. The first one (general proteins) is obtained from the `ProteinMPNN` datasets (see 2.2.1). The second one (specific proteins) is a selection of specific proteins from the PDB.

#### 3.1.1 General proteins

We use the same data as was used in the original `ProteinMPNN` paper ^1^, also keeping their train, validation and test split (see 2.2.1). Based on the training data, we generate our preference datasets for alignment (see 3.2). 98 randomly selected proteins of the validation data are used to perform the trade-off analysis (see 3.4).

#### 3.1.2 Specific proteins

We use four monomer (single amino-acid chain) protein structures (IDs: 2R4U, 5BX2, 3X2G, 2EID) as well as two homo-dimers (two some amino-acid chains) protein structures (IDs: 4RGD, 3BBE) from the PDB in the hyper-parameter assessment process. We call these the specific validation proteins.

In contrast, the specific test proteins are monomers and homooligomers that we actually want to apply our method to. We do not ensure that these are not present in the training or validation set. Their IDs are: 4KW4, 8A6G, 1UBQ, 2QMT, 6QJI, 1M40, 5O75, 1EUM, 1QAW, 4GYT, 6EHB.

### 3.2 Alignment

The alignment process is meant to shape the responses of a model in a way, so that they better align with our preferences. In the Natural Language Processing (NLP) setting the prompt might be a question to which the LLM is supposed to respond with a helpful and harmless answer. In contrast, in our protein design setting the prompt is a protein backbone, to which the tuned `ProteinMPNN` is supposed to responds with a particularly low immune-visible corresponding AA sequence.

The basic hypotheses of this work is that the same DPO process as used for the NLP setting can also be used in the protein design setting.

As base model for the alignment we use the `v 48 020` version of `ProteinMPNN` as described in 2.2. As seen in Subsection 2.3, DPO tuning requires the creation of a preference pair dataset.

#### 3.2.1 Preference dataset

This dataset is generated in the following way. For every backbone example in the general training data (see 3.1), we let the current alignment state of `CAPE-MPNN` generate two candidate designs. We require the network to generate all AAs in the sequence. These two candidates are then assessed by our bioinformatics assessment pipeline (see 3.2.2) and one is selected as the preferred example. In addition, we also use the original `ProteinMPNN` weights to calculate the likelihood of the two examples being generated as responses to the backbone. With this preference dataset, the model is then aligned for two epochs. Then a new preference dataset is generated using the latest aligned version of `CAPE-MPNN`. This is in contrast to the original DPO paper [9], which uses the original model to generate the preference examples, we use the tuned one. The reasoning behind this is that while in language tasks an answer is typically either acceptable or not, in our task we want to move the model further and further away from the original outputs to gradually reduce immune-visibility. We think that, therefore, it is necessary to generate preference examples that reflect the current state of tuning. However, the *π*_*ref*_ used in the loss in equation 1 remains the one of the original `ProteinMPNN` base model.

#### 3.2.2 Assessing preference of designs

Although DPO is based on RLHF, which has “human feedback” in its name, we replace this by feedback from bioinformatics methods that assess the immune-visibility of generated sequences. Less visible ones are preferred.

In this first iteration of `ProteinMPNN` we only consider visibility via the MHC-I pathway. State of the art MHC-I presentation prediction methods like *netMHCpan 4*.*1* [21] can be quite slow. So we constructed position weight matrix (PWM) classifiers based on the predictions of *netMHCpan 4*.*1* [21] on whether peptides were presented or not. Each PWM has one row per AA and one column per position in the peptide. The values are the log-likelihood probably to find this row’s AA in the column’s position in peptides being presented by the MHC-I pathway (details see 2.1. [22]).

### 3.3 Assessment criteria

Whenever a 3D structure to a designed protein is needed, we used ColabFold wich is based on AlphaFold [10, 23, 24] to predict this.

#### Visibility

We define visibility as in `Box1`. However, we will typically not use the absolute number of this, but rather compare the visibility of the design with the visibility of the original sequence in the dataset. So if the designed protein has a visibility of 18, while the original one had 36, we will use 0.5.

#### Sequence recovery

Similar to the original paper [1], we also look at the percentage of AAs that are the same in the natural protein as in the generated sequence.

#### TM-score

A common method to compare the structural similarity between two proteins is the TM-score. This can be calculated with *TMalign* [25]. The TM-score can range between zero and one. While scores below 0.2 are associated with random, unrelated proteins, scores that are higher than 0.5 point to the same fold in SCOP/CATH ^2^.

### 3.4 Trade-off analysis

To analyse the trade-off between visibility reduction and quality of the generated proteins, we conduct a DPO hyper-parameter search for the reduction of immune-visibility in a hypothetical patient with the following MHC-I alleles: HLA-A*02:01, HLA-A*24:02, HLA-B*07:02, HLA-B*39:01, HLA-C*07:01 and HLA-C*16:01). Each run was performed with a different randomly sampled hyper-parameter set (see **Table 2**). The trade-off analysis is then based on twenty training runs over twenty epochs each. Of these 20 hyper-parameter runs, three failed due to stability issues. These seem to have been the result of too high *β* values. Of the succeeding runs, we selected the checkpoints after epoch 2, 10 and 20. This resulted in a total of 17×3 checkpoints to compare. For each **specific validation protein**, we let each of these checkpoints generate 3 candidate protein designs (using a temperature of 0.1). Since we observed that some of the checkpoints would output only constant chains, we remove all candidate sequences which include less than 15 different AA types. We then use ColabFold [10, 23, 24] to generate 3D structures for each of the remaining candidate designs (unsuccessful design get assigned a TM-score of zero). In case there are candidates with TM-scores above 0.9, we select the one with the minimum visibility from those. Otherwise, we select the candidate that has the maximum TM-score. For the analyse of the trade-off between visibility reduction and quality, we use visibility reduction over general validation proteins as proxy for visibility reduction and TM-score on the predicted specific validation proteins as proxy for quality.

**Table 2:** CAPE-MPNN DPO hyper-parameters.

| DPO hyper-parameter | Distribution | Value |  |
| --- | --- | --- | --- |
|  |  | 458340e4 | e32b8ed0 |
| beta | log uniform(0.001, 0.2) | 3.43e-2 | 4.48e-3 |
| sample temperature | uniform(0.01, 1.0) | 0.759 | 0.500 |
| learning rate | log uniform(1e-7, 1e-3) | 2.86e-6 | 4.62e-7 |

### 3.5 Specific protein analysis

Finally, we selected two promising models identified in the trade-off analysis (`458340e4` and `e32b8ed0`, see **Table 2**) to analyse their performance on some more specific test backbones. Again, we analyse the results with regards to TM-score and visibility. However, we now present numbers for specific designs instead of averages.

## 4 Results and Discussion

This section first presents the results of the trade-off analysis that was carried out using validation set backbone structures. Afterwards, we look into the performance of two models on specific backbone structures in more detail.

### 4.1 Trade-off analysis

To analyse the dependence of quality of generated sequences (as measured by TM-score) and visibility modification achieved, we tuned `CAPE-MPNN` checkpoints with various hyper-parameters (see **Table 2**).

**Figure 1** that plots the visibility modification vs. the quality of the designs shows that there exists a trade-off between those two objectives. Both plots in the figure show that the `ProteinMPNN` base model (‘v 48 020’) designs in red are high quality but with similar visibility to the original proteins. In comparison, the `CAPE-MPNN` designs experience a trade-off between quality and visibility, dependent on the hyper-parameters used to train them.

**Figure 1:**
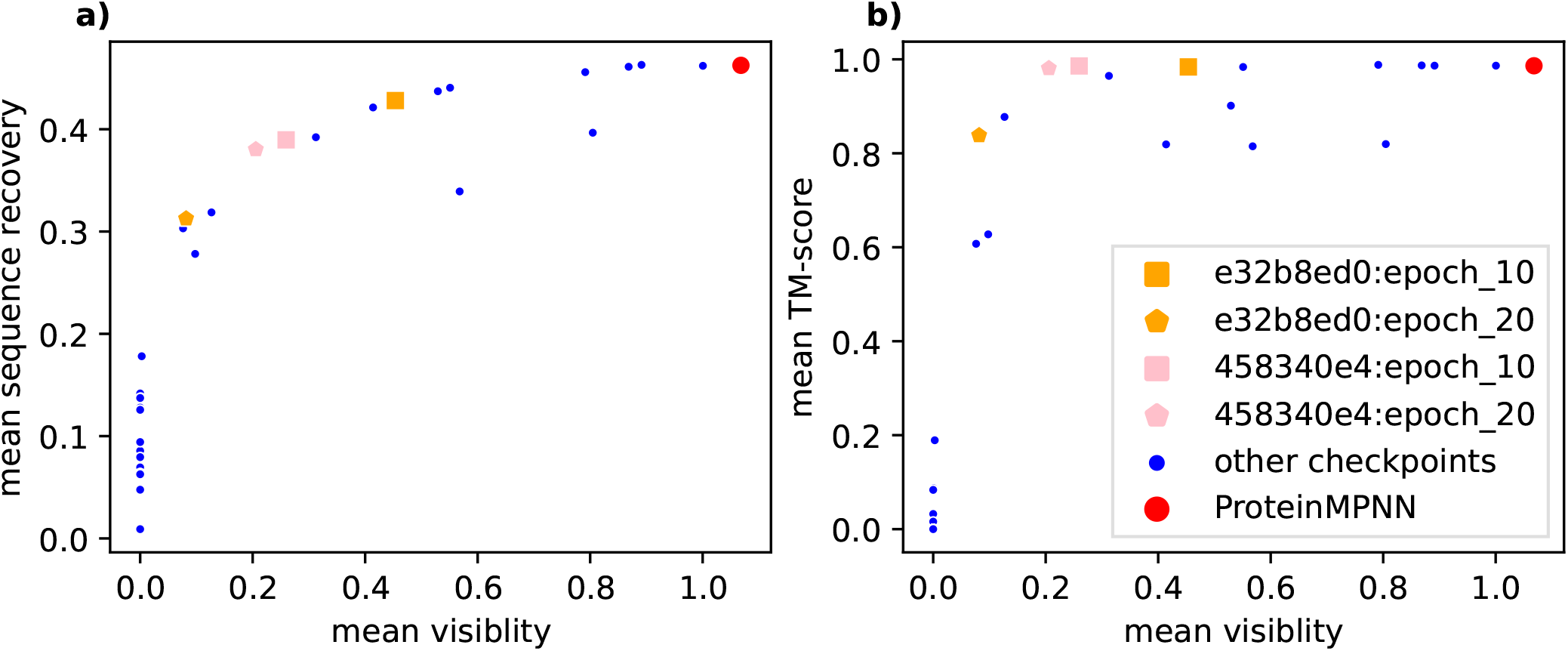
Trade-off between visibility and structural validity on validation data: Each data-point in these two plots represents a checkpoint. The red one is the original `ProteinMPNN` network. The blue ones represent checkpoints that were created during the hyper-parameter search (after 2, 10 and 20 epochs). The orange and pink ones are for two selected hyper-parameter combinations. Both plots show the mean relative visibility to MHC-I over the general validation proteins on the x-axis. (a) shows the mean sequence recovery over the validation proteins while (b) shows the mean TM-score over the specific validation protein designs on the y-axis

**Figure 1a** shows that the reduction in sequence recovery that is linked with a reduction in visibility seems to be quite steady and there seems to be a narrow band of attainable recovery values for a given visibility. This points to most checkpoints being “pareto optimal” with respect to the visibility/quality trade-off -independent of their actual hyper-parameters.

In comparison, **Figure 1b** depicts that the predicted TM-scores stay quite high before suddenly falling off a cliff at a level of roughly 20% the original visibility. We also observe a wider band of mean TM-scores for a given mean visibility. This might have to do with the relative few specific designs we used for the calculation of the mean TM-score. Any failed design will attract a TM-score of zero and so have a large negative impact on the value.

To identify which hyper-parameters determine the eventual position of a model on this trade-off frontier, **Figure 2** depicts the dependence of TM-score and visibility on the three varying hyper-parameters (**Table 2**) after 2 epochs of tuning. We found that high beta values would lead to unstable training (not depicted in the figure). Learning rate (lr), seems to have an influence on the eventually achieved quality and visibility modification. Unsurprisingly, high learning rates seem to “destroy” the `ProteinMPNN` weights and make the model output gibberish. In contrast, very low learning rates, do not modify the weights at all. Raising learning rate then first seems to affect visibility in a positive way before leading to reductions in TM-score. The temperature used to sample the preference examples, seems to have little influence on the eventual outcome.

**Figure 2:**
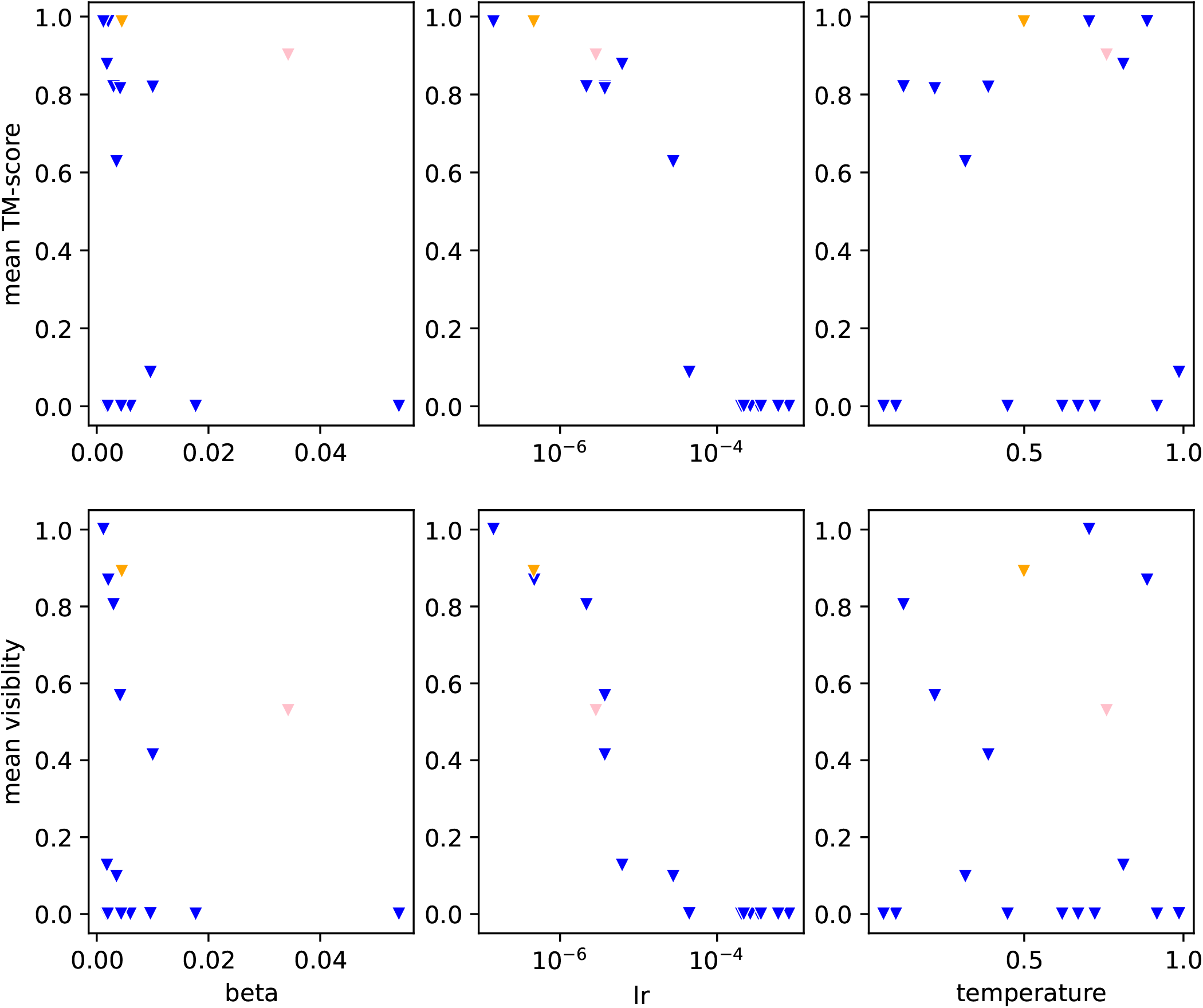
Influence of DPO-hyperparameters on the quality vs. visibility modification trade-off: Each triangle in the plots corresponds to a single checkpoint (after epoch 2 of tuning). The colors are the same as in **Figure 1**. Each column of the figure shows the influence of a variation in a DPO-hyperparameter. The first row depicts the influence on the mean TM-score of the designs for the specific validation backbones. The second row depicts the influence on the mean visibility of the designs for the general validation backbones.

### 4.2 Specific protein analysis

With regards to specific proteins, we observe a broad range of performance (**Figure 3**). There are some proteins, for which the results are quite encouraging. In particular, we find that `458340e4:epoch 20` and `e32b8ed0:epoch 20` designed candidates for GB1 of *Staphylococcus aureus* (PDB: 2QMT) that have a predicted visibility of zero while the predicted TM-score retains a high value of above 90%. In contrast, we see that the `ProteinMPNN` base model in **Figure 3a** generated candidates, that all had visibilities above 50% of the original visibility. We want to point out that this is a fair comparison, as we also created three candidate designs for each protein for the base model.

**Figure 3:**
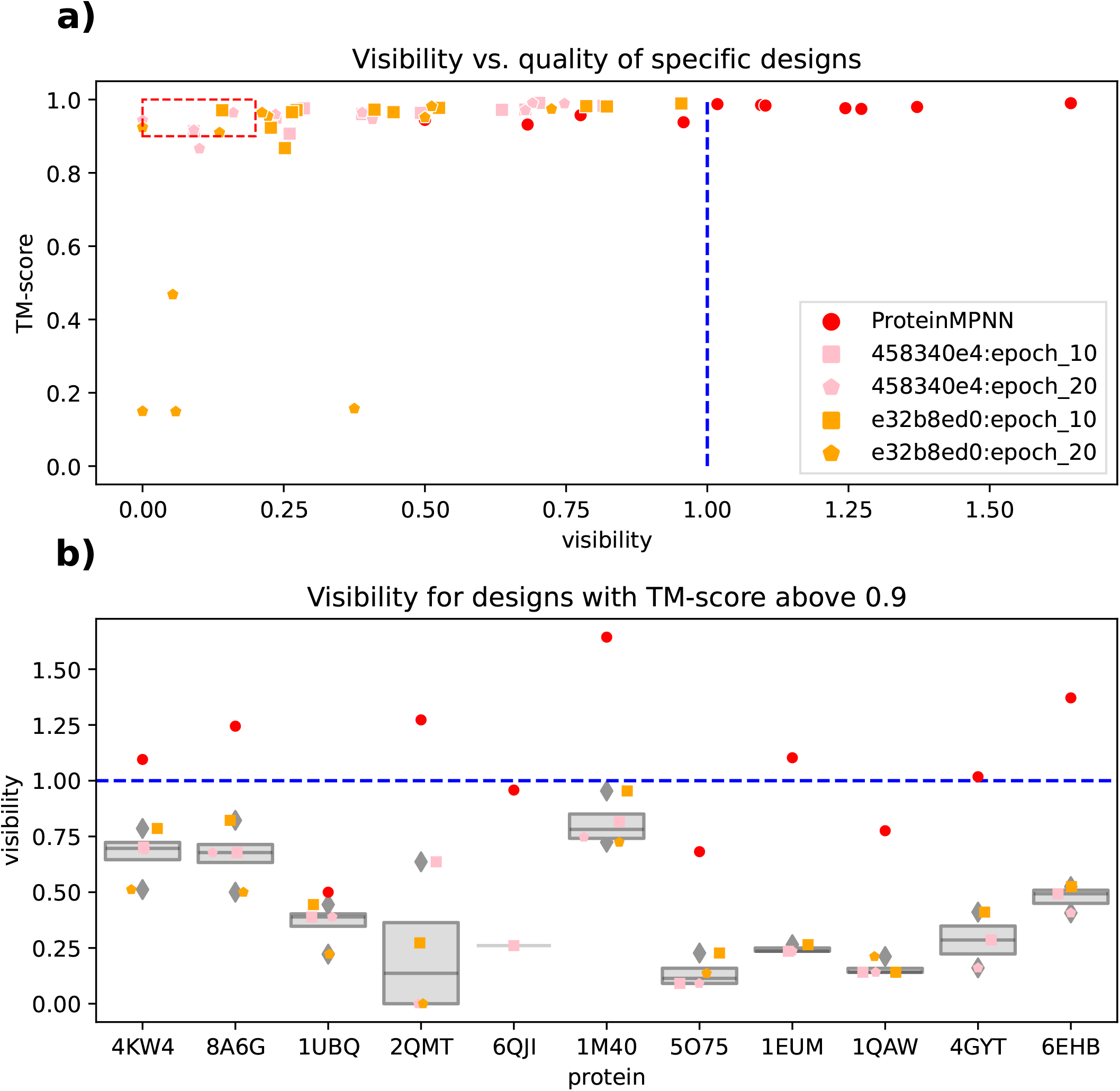
A broad performance range: This figure looks into specific designs. In plot **a)** the quality of generated specific designs (as measured by TM-score on the y-axis) gets compared to the relative visibility (x-axis) for two DPO hyper-parameter selections after two epochs of tuning. Each point represents a design for a specific backbone using the checkpoint. We find that most designs have high TM-scores above 0.9. Not all of them are less visible than the original sequence (visibility *<* 1.0) and we also see that the `ProteinMPNN` designs in red tend to have far higher visibility. Furthermore, some designs also only show a maximum of 20% the number of visible peptides in the designs (red dashed box). In plot **b)** we then look into the distribution of visibilities of designs for specific backbones. These are subset of designs from plot a) that satisfy the condition that their TM-score is above 0.9. The dashed blue lines represent the visibility of the PDB proteins.

## 5 Conclusion

The renewed interest in protein design kindled by advances in ML has led to steady advances in this area. To use these de novo proteins as therapeutics in actual patients, prevention of unwanted immune-reactions is paramount. In this paper we analyse the potential of using DPO to tune the state of the art (SOTA) protein design model `ProteinMPNN` to generate less MHC-I immune-visible proteins. We explore the trade-off between reducing visibility and quality of the generated designs. The findings point to the ability of DPO to effectively integrate a deimmunization goal into the `ProteinMPNN` protein design process. However, it also demonstrates it limitations, as visibility cannot be reduced to zero in the vast majority of cases.

## Acronyms

AA: amino acid
Ab: antibody
AR: auto-regressive
CTL: Cytotoxic T-lymphocyte
DPO: Direct Preference Optimization
GAN: Generative Adversarial Network
LLM: large language model
MD: Molecular Dynamics
MHC-II: MHC Class II
MHC-I: MHC Class I
ML: machine learning
MPNN: message passing neural network
NLP: Natural Language Processing
PPO: Proximal Policy Optimization
PWM: position weight matrix
RBF: radial basis function
RL: reinforcement learning
RLHF: reinforcement learning from human feedback
RNA: ribonucleic acid
SOTA: state of the art
VAE: Variational Autoencoder

## 6 Acknowledgements

This work was supported by the United Kingdom Research and Innovation (grant EP/S02431X/1), UKRI Centre for Doctoral Training in Biomedical AI at the University of Edinburgh, School of Informatics. For the purpose of open access, the author has applied a creative commons attribution (CC BY) licence to any author accepted manuscript version arising. This project was supported by the Royal Academy of Engineering and the Office of the Chief Science Adviser for National Security under the UK Intelligence Community Postdoctoral Research Fellowship programme.

J.A. Alfaro was supported by (i) United Kingdom Research and Innovation (grant EP/S02431X/1), (ii) the project ‘International Centre for Cancer Vaccine Science’ which is carried out within the International Agendas Program of the Foundation for Polish Science, cofinanced by the European Union under the European Regional Development Fund. The authors would like to thank ‘CI-TASK, Gdansk’, and the ‘PL-Grid Infrastructure, Poland’ for providing their hardware and software resources. Dr. Rajan was supported by the KATY Consortium H2020-SCI-FA-DTS-2020-1.

## Footnotes

1 https://github.com/dauparas/ProteinMPNN/tree/main/training

2 https://seq2fun.dcmb.med.umich.edu/TM-align/

## Notes

### Competing Interest Statement

The authors have declared no competing interest.

